# Engineering a Controlled Cardiac Multilineage Co-Differentiation Process Using Statistical Design of Experiments

**DOI:** 10.1101/2025.03.18.643864

**Authors:** Hirokazu Akiyama, Yosuke Katayama, Kazunori Shimizu, Hiroyuki Honda

**Author notes:** Co-correspondence: Hirokazu Akiyama, Hiroyuki Honda.

## Abstract

**Background:** The heart is a complex organ composed of diverse cell types, whose interplay is crucial for development, tissue homeostasis, and disease progression. A potential approach to recapitulate this heterotypic multicellular environment is the co-differentiation of induced pluripotent stem cells (iPSCs) into multiple cardiac cell types, facilitating the unlocking of their full potential for regenerative medicine and drug development. However, the inherent complexity of co-differentiation, where multiple differentiation factors simultaneously influence the induction of multiple cell types, presents a significant challenge in achieving high controllability over the process toward the desired outcome. Thus, a robust strategy is essential for engineering a controlled co-differentiation process with broader applicability.

**Methods:** Given the importance of heterotypic cellular proportions in facilitating proper interactions, we present a new strategy to engineer a controlled cardiac co-differentiation process from iPSCs using statistical design of experiments to simultaneously generate cardiomyocytes, mural cells, and endothelial cells, which are major constituents of the heart.

**Results:** We divided the process into two stages: progenitor cell induction and the subsequent trilineage co-differentiation, allowing for stage-specific optimization. Given that the performance of progenitor cell induction may critically influence the overall process performance, we carefully optimized activin A and CHIR-99021 using the sequential design of experiments to achieve approximately 95% induction efficiency of KDR^+^/PDGFR-α^+^ cardiogenic mesoderm cells from iPSCs with minimal batch-to-batch variability. In the trilineage co-differentiation stage, we developed unique multi-response models to delineate trilineage co-differentiation ratios within a defined parameter space of WNT signal inhibitor and vascular endothelial growth factor. This enabled the identification of potential conditions that steer co-differentiation toward desired cellular constitutions, a critical factor of effective cellular interplays. Repeated trilineage co-differentiation experiments confirmed the high process controllability, with a close match between actual and predicted differentiation ratios. Furthermore, cardiomyocytes from trilineage co-differentiation exhibited a more mature sarcomere gene expression profile than those from monolineage differentiation.

**Conclusions:** These results highlight the effectiveness of our strategy for engineering a stem cell co-differentiation process and its applicability where multicellular interactions are crucial.

## Background

The heart is the first solid organ to form during embryonic development and consists of diverse cell types arising from distinct precursor populations, namely cardiogenic mesoderm, proepicardium, and cardiac neural crest [1,2]. Among them, the cardiogenic mesoderm gives rise to the major constituents of the heart, including cardiomyocytes (CMs), endothelial cells (ECs), and mural cells (MCs), which interact and play essential roles in the structural and functional development of the hearts [1]. The interplay among multiple cell types through physical contact and/or paracrine signaling is crucial not only for heart development but also for tissue homeostasis and disease progression [3,4]. Therefore, bioengineering strategies to recapitulate the heterotypic multicellular environment are indispensable for unlocking the authentic potential of cardiac cells for applications such as regenerative medicine and drug development as well as for studying cardiac biology.

Among pluripotent stem cell (PSC) derivatives, CMs have been one of the most extensively studied cell types, with applications in regenerative medicine [5,6] and drug development [7–9] gradually expanding. To recapitulate a multicellular environment involving PSC-derived CMs, the co-culture approach, where different cell types are prepared separately and then combined, has been the de facto standard and has demonstrated its effectiveness in modulating CM properties [10–14]. Alternatively, co-differentiation, in which multiple cell types are simultaneously induced from stem cells by recapitulating developmental processes, represents another possible approach [15,16]. Co-differentiation offers some advantages over co-culture in biological and technical aspects; it more closely mimics the natural cell-cell interactions between multiple heterotypic cells during development and eliminates the need to establish multiple cell preparation protocols or laboriously preparing them independently. However, owing to the inherent complexity of co-differentiation, where multiple differentiation factors simultaneously influence the induction of multiple cell types, the development of a controlled process that achieves the desired outcome remains highly challenging. Specifically, given the importance of heterotypic cellular proportions in facilitating proper interactions, as observed in the aforementioned co-culture studies, achieving high controllability over the differentiation ratio of multiple cell types is a key focus. Therefore, a robust strategy is essential for engineering a controlled co-differentiation process with broader applicability.

Design of experiments (DoE) is a powerful process engineering approach that investigates relationships between multiple factors and outcomes through statistical modeling [17–20]. While extensively used across various industrial fields, its application in scientific research appears to be less prevalent despite its substantial utility. It is highly useful not only in factor screening and optimization for a single response but also in multi-response investigation, due to its ability to efficiently reveal multiple response behaviors within a defined multidimensional parameter space. A notable example of multi-response investigation using the DoE approach is a recent study by the Zandstra group [20], where they successfully established a T cell generation process from hematopoietic stem and progenitor cells by optimizing multiple cytokines to maximize multiple distinct cell populations in a stage-dependent manner. However, to the best of our knowledge, no studies have demonstrated the application of DoE in engineering a co-differentiation process to control differentiation ratios among distinct lineage cells. Thus, we explored the potential of the DoE for the development of a controlled cardiac multilineage co-differentiation process.

In this study, we present a new strategy to develop a co-differentiation process to generate CMs, MCs, and ECs with a controlled differentiation ratio from iPSCs using the DoE approach. We divided the process into two stages: cardiogenic mesoderm induction and the subsequent trilineage co-differentiation, allowing for stage-specific optimization. Since the performance of the initial stage differentiation significantly influences the overall process performance, we conducted sequential DoE experiments to carefully select and optimize factors for efficient, stable, and reproducible cardiogenic mesoderm induction. In the later core stage, we developed multi-response models to reveal the differentiation ratios of these trilineage cells within a defined parameter space, allowing us to select the potential conditions that steer differentiation toward the targeted cellular ratios. We then tested the controllability of the co-differentiation process for the selected conditions by assessing whether the actual and predicted differentiation ratios closely matched. Finally, we compared cardiac maturation-associated gene expression in CMs from tri and monolineages differentiation to evaluate the impact of co-differentiation on CM properties.

## Methods

### Human iPSC maintenance

The 253G1 hiPSC line [21], obtained from the RIKEN BioResource Research Center, was used in this study. The iPSCs were adapted to and maintained in Essential 8 medium (E8; Thermo Fisher Scientific, Waltham, MA, USA, Cat. # A1517001) on dishes coated with VTN-N (Thermo Fisher Scientific, Cat. # A14700), with daily medium replacement. Clump passaging was performed every 3–4 days using 0.5 mM ethylenediaminetetraacetic acid solution, with a split ratio of 1:8.

### Factor evaluation for cardiogenic mesoderm induction

iPSCs were detached and dissociated using accutase (Nacalai Tesque, Kyoto, Japan, Cat. # 12679-54) and seeded into a 48-well plate (Greiner Bio-One, Kremsmunster, Austria, Cat. # 677180) pre-coated with Geltrex (1:50 dilution; Thermo Fisher Scientific, Cat. # A1413302) at 20,000–25,000 cells/cm^2^ in E8 medium supplemented with 10 μM Y-27632 (Nacalai Tesque, Cat. # 18188094). The cells were expanded for three days before differentiation.

To assess the effects of activin A (Act; R&D Systems, Minneapolis, MN, USA, Cat. # 338-AC), bone morphogenetic protein-4 (BMP; PeproTech, Cranbury, NJ, USA, Cat. # 120-05ET), and CHIR-99021 (CHIR; Cayman Chemical, Ann Arbor, MI, USA, Cat # 13122) during the first 24 h of differentiation on cardiogenic mesoderm induction, a full factorial design experiment was conducted. Differentiation media with varying concentrations of the three factors were formulated according to the design matrix in Table S1. The media were based on RPMI 1640 (Nacalai Tesque, Cat. # 30264-85) supplemented with 2% B27 minus insulin (Thermo Fisher Scientific, Cat # A1895601), hereafter referred to as RPMI/B27 (-ins). A liquid handler (OT-2; Opentrons, Long Island City, NY, USA) was employed for precise media formulation in all DoE experiments throughout this study. When the iPSCs reached approximately 90% confluence, the medium was replaced with the differentiation media, defining this day as day 0. Exactly 24 h later, the media were replaced with RPMI/B27 (-ins). Cells were collected for marker expression analysis on days 3 and 4 for each condition to accommodate the transient, condition-dependent peak timing of marker expression. Percentages of three cardiogenic mesoderm populations (KDR^+^, PDGFR-α^+^, and KDR^+^/PDGFR-α^+^) and one non-cardiogenic mesoderm population (KDR^−^/PDGFR-α^−^) were measured using flow cytometry, as described later. For each condition, the highest value observed on either day for cardiogenic mesoderm or the lowest value for non-cardiogenic mesoderm was used to estimate parameter coefficients in the following regression model:

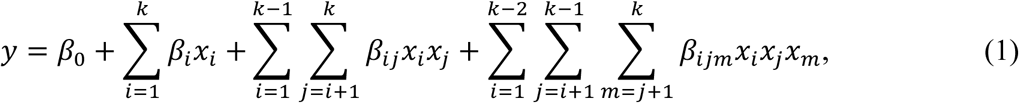

where *β*_0_ represents the intercept; *x_i_*, *x_j_*, and *x_m_* represent the factor-scaled levels shown in Table S1; *β_i_*, *β_ij_*, and *β_ijm_* are the coefficients for the main effects, second-order interactions, and third-order interactions; and *y* represents percentage of the population of interest.

### Optimization of cardiogenic mesoderm induction

To optimize the concentrations of CHIR and Act, response surface methodology based on a central composite design was conducted for the KDR^+^/PDGFR-α^+^ population. The experiment was conducted twice with slightly different concentration ranges according to the design matrix in Table S2, using the same experimental procedure as in the factor evaluation. Pooled data from both experiments were used to estimate parameter coefficients in the following quadratic model:

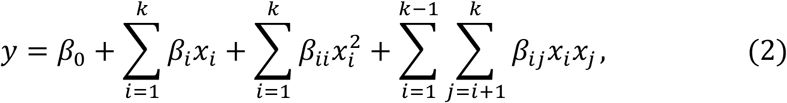

where *x_i_* and *x_j_* represent the factor-scaled levels, recoded such that −1 and +1 correspond to the lowest and highest concentrations across the two experiments; *β_i_*, *β_ii_*, and *β_ij_* are the coefficients for the main effects, quadratic effects, and second-order interactions; and other variables remain as described earlier.

The optimized condition was tested by measuring the percentage of KDR^+^/PDGFR-α^+^ cells using the same culture scheme. The resultant data were then compared with the model’s prediction, which incorporated random errors based on an error distribution defined by the root mean square error.

Upon changing a CHIR lot, a dose-response assessment was conducted, as the optimal concentration for cardiac differentiation has been reported to be lot-dependent [22]. This may be partially due to minor weighing errors during powder vialing at the manufacture and/or reconstitution in the laboratory. Differentiation was initiated in a 48-well plate with CHIR at −1 to + 1 μM of the initially optimized concentration, in the presence of Act at the optimized concentration, followed by the same culture scheme. On day 3, cells were harvested and analyzed using flow cytometry to determine the percentages of the KDR^+^/PDGFR-α^+^ cells, using the procedure described later. The CHIR concentration that yielded the highest KDR^+^/PDGFR-α^+^ cell induction was selected for experiments with the new lot to ensure experimental reproducibility throughout this study.

### Optimization for cardiac trilineage co-differentiation

To predict the differentiation ratios of CMs, MCs, and ECs during co-differentiation in response to varying WNT signaling inhibitor (WNTi) and vascular endothelial growth factor (VEGF; Qkine, Cambridge, UK, Cat # QK048), a response surface methodology was performed. Herein, WNTi consisted of a 1:1 molecular ratio mixture of IWP-2 (Cayman Chemical, Cat. # 13951) and XAV-939 (Selleck Chemicals, Houston, TX, USA, Cat. # S1180) with the indicated concentration representing the total concentration of both compounds. Differentiation was initiated with the optimized CHIR and Act concentrations using a 48-well plate. On day 2, the media were replaced with RPMI/B27 (-ins) containing varying concentrations of the two factors in accordance with the design matrix in Table S3. From day 4 onward, VEGF was retained, while WNTi was excluded, with media exchanged every 2–3 days. On day 10, the cells were harvested and subjected to flow cytometry analysis to measure the percentages of CMs (cTNT^+^), MCs (PDGFR-β^+^/cTNT^−^/VE-cad^−^), and ECs (VE-cad^+^/cTNT^−^/PDGFR-β^−^) using the procedure described later. The results were used to estimate parameter coefficients in the quadratic model defined in Equation (2), where *x_i_* and *x_j_* represent the factor-scaled levels shown in Table S3, while other variables remain as described earlier.

The controllability of differentiation ratios for the selected conditions was tested using the same differentiation scheme. In parallel, monolineage differentiation was performed, where differentiation was initiated under the same condition but continued with 10 μM WNTi from days 2 to 4 to exclusively generate CMs. The results of co-differentiation were then compared with the model’s predictions, which incorporated random errors based on an error distribution defined by root mean square error.

### Prolonged aggregation culture

On day 10 of differentiation, mono and trilineage differentiation cells were harvested and dissociated using the AscleStem Cardiomyocyte Dissociation Solution (Nacalai Tesque, Cat. # 17080-40) according to the manufacturer’s protocol. The cells were then suspended in α-MEM (Nacalai Tesque, Cat. # 21444-05) supplemented with 10% fetal bovine serum (Sigma-Aldrich, St. Louis, MO, USA, Cat. # 172012), 1% penicillin-streptomycin (Fujifilm Wako Pure Chemical, Osaka, Japan, Cat. # 168-23191), and 10 μM Y-27632, with or without 50 ng/mL VEGF for tri or monolineage cells, respectively. The cells were then seeded into an Elplasia 96-well plate (Corning, Corning, NY, USA, Cat. # 4442) with low attachment microcavities at 316,000 cells/well, equivalent to 4,000 cells per cavity. Immediately thereafter, the plate was centrifuged at 100 × *g* for 3 min to facilitate aggregate formation. After 48 h, the medium was replaced with RPMI 1640 supplemented with 2% B27 containing insulin (Thermo Fisher Scientific, Cat. # 17504044) and 1% penicillin-streptomycin, hereafter referred to as RPMI/B27 (+ins), with or without 50 ng/mL VEGF for tri or monolineage cells, respectively. The medium was replaced every 2–3 days. On day 40, equivalent to day 30 of the aggregation culture, cells were subjected to gene expression analysis using the procedure described later.

### Flow cytometry analysis

Cells were harvested using TrypLE Express (Thermo Fisher Scientific, Cat. # 12604013). For cardiogenic mesoderm analysis, cells were stained at 4°C for 30 min with fluorophore-conjugated antibodies targeting surface proteins, followed by washing with phosphate-buffered saline containing 2% fetal bovine serum and 0.5 mM ethylenediaminetetraacetic acid, hereafter referred to as FCM buffer. For CM, MC, and EC analysis, cells were stained with the LIVE/DEAD Fixable Violet Dead Cell Stain Kit (Thermo Fisher Scientific, Cat. # L34963) following the manufacturer’s protocol. Thereafter, cells were stained at 4°C for 30 min with fluorophore-conjugated antibodies targeting surface proteins. Then, cells were fixed and permeabilized using the Foxp3/Transcription Factor Staining Buffer Set (Thermo Fisher Scientific, Cat. # 00-5523-00) in accordance with the manufacturer’s instructions. Following this, cells were stained at 4°C for 30 min with a fluorophore-conjugated antibody targeting an intracellular protein, followed by washing with FCM buffer. The stained samples were analyzed using the BD FACSCanto II flow cytometer (Becton Dickinson, Franklin Lakes, NJ, USA) equipped with a high-throughput sampler, after which the data were analyzed using FlowJo software (version 10.8 or 10.9; Becton Dickinson). The antibodies used in this analysis are listed in Table S4.

### Gene expression analysis

RNA was extracted using the NucleoSpin RNA Plus XS kit (Macherey-Nagel, Dueren, Germany, Cat. # 740990) following the manufacturer’s protocol. For each sample, 100–150 ng of RNA was reverse-transcribed into cDNA using the ReverTra Ace qPCR RT Master Mix with gDNA Remover (Toyobo, Osaka, Japan, Cat. # FSQ-301) at 37°C for 15 min, followed by 50°C for 5 min. Quantitative PCR (qPCR) was conducted using 1.5 ng of cDNA as a template with the Thunderbird SYBR qPCR Mix (Toyobo, Cat. # QPS-201) on the StepOne Real-Time PCR System (Thermo Fisher Scientific). The qPCR conditions consisted of an initial denaturation step at 95°C for 60 s, followed by 40 cycles of denaturation at 95°C for 15 s, annealing at 59°C for 15 s, and extension at 72°C for 30 s. Relative expression levels were calculated using the ΔΔCt method, with *RPL37A* as an endogenous gene control. The primers used in this study are listed in Table S5.

### Statistical analysis

Parameter coefficients of the models defined in Equations (1) and (2) were estimated using regression analysis with the least squares method in JMP Pro (version 16, via an institutional license; JMP Statistical Discovery, Cary, NC, USA), and their statistical significance was assessed using the *F*-test, with *p*-values adjusted using the Benjamini-Hochberg (BH) method. A single pairwise comparison was performed using a *t*-test, while multiple pairwise comparisons were performed using one-way analysis of variance (ANOVA), followed by Tukey’s post hoc tests unless otherwise noted. Multivariate comparisons were conducted using multivariate ANOVA. Any *p*-values less than 0.05 were considered statistically significant.

## Results

### Activin A, not BMP-4, promotes cardiogenic mesoderm induction with suboptimal CHIR-99021

The overall process outcome is highly dependent on the performance of the initial differentiation from iPSCs to cardiogenic mesoderm, necessitating robust optimization to ensure efficient and reproducible differentiation through precise modulation of signaling pathways. WNT signaling activation via CHIR-99021 (CHIR), a glycogen synthase kinase-3β inhibitor, is the most common approach to initiate cardiac differentiation [23–25]. Bone morphogenetic protein-4 (BMP) and activin A (Act) have also been used to precisely mimic primitive streak development [26–28]. Additionally, CHIR has been combined with both BMP and Act [13,16], or with either factor [22,29,30]. Yet, the optimal combination remains inconclusive, prompting us to investigate their effects on cardiogenic mesoderm induction.

For this purpose, we introduced these three factors during the first 24 h of differentiation from monolayer hiPSCs using a two-level full factorial design (FFD) (Fig. 1a, b), which allowed simultaneous evaluation of the main effects and interactions. In this FFD experiment, while Act and BMP were set at 0 and 10 ng/mL for the low and high levels, CHIR was set at 8 and 10 μM, respectively (Table S1). This concentration setup was specifically designed to assess the effects of BMP and Act in the presence of suboptimal CHIR. Furthermore, we used an automated liquid handler for media formulation throughout all DoE-based experiments to obtain high-precision data throughout this study. For the evaluation of cardiogenic mesoderm induction, we employed two surface markers, KDR and PDGFR-α, widely used in combination [27,31] or sometimes individually [32,33] for this purpose. As shown in Fig. 1c, factorial analysis revealed very significant positive main effects of Act and CHIR on three cardiogenic mesoderm populations (KDR^+^, PDGFR-α^+^, and KDR^+^/PDGFR-α^+^). In contrast, both factors showed very significant negative effects on the non-cardiogenic mesoderm (KDR^−^/PDGFR-α^−^), which was consistent with their opposite influence on cardiogenic mesoderm. No significant effects associated with BMP were observed for any population. These findings indicate that Act, but not BMP, supports cardiogenic mesoderm induction in the presence of suboptimal CHIR.

**Fig. 1.**
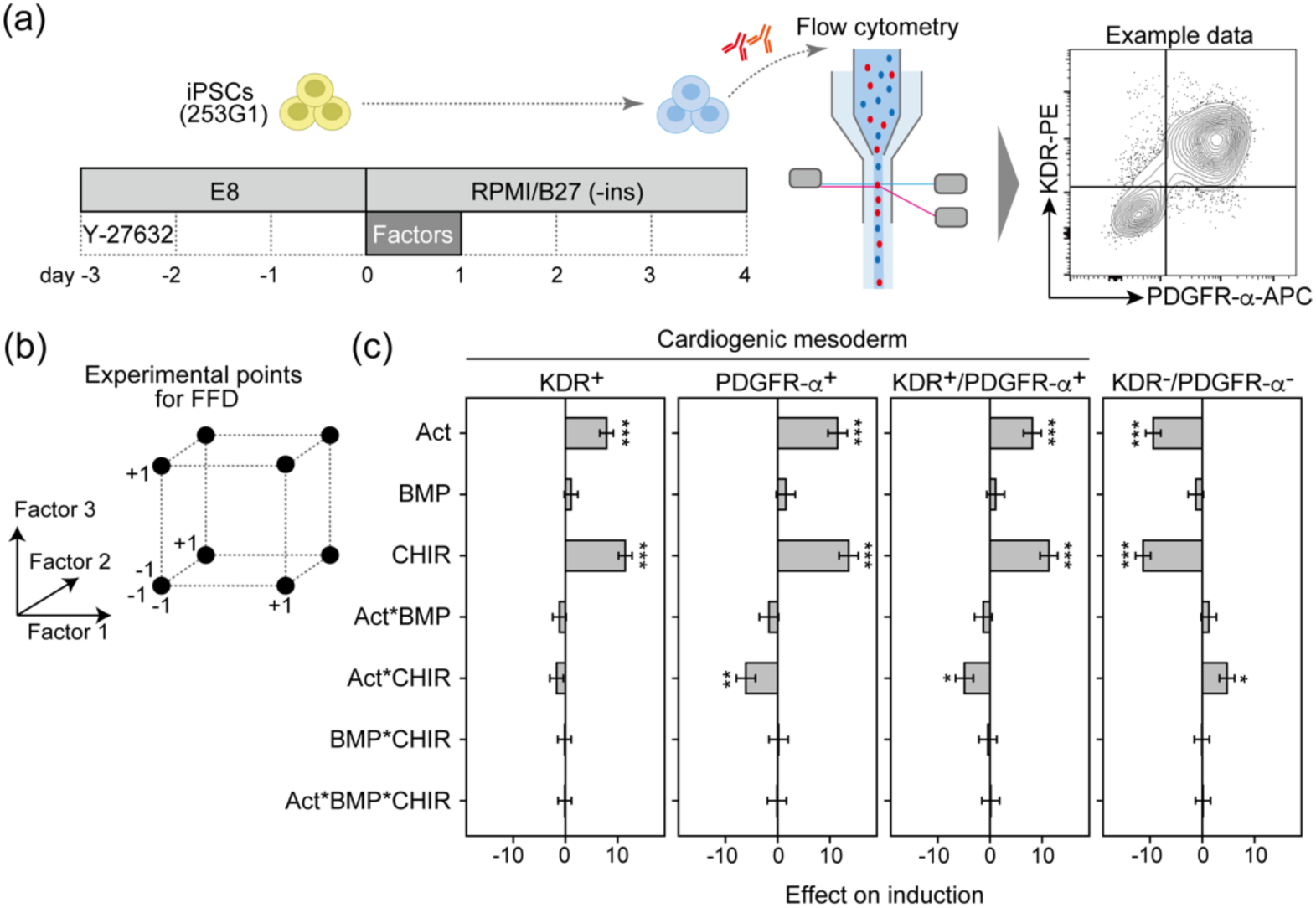
Factor evaluation for cardiogenic mesoderm induction. (a) Schematic representation of the experimental workflow for evaluating the effects of activin A (Act), BMP-4 (BMP), and CHIR-99021 (CHIR) on cardiogenic mesoderm induction. These factors were introduced during the first 24 h of differentiation at the concentrations specified in Table S1, based on a full factorial design (FFD). (b) Geometric representation of experimental points in the FFD with three factors. The scaled values “−1” and “+1” represent the low and high concentrations, respectively. (c) Results of factorial analysis on induction efficiencies of cardiogenic mesoderm populations (KDR^+^, PDGFR-α^+^, and KDR^+^/PDGFR-α^+^) and a non-cardiogenic mesoderm population (KDR^−^/PDGFR-α^−^). The effects are shown as the values of the corresponding coefficients in Equation (1), along with standard errors (SEs). Statistical significance was assessed using an *F*-test, followed by *p*-value adjustment using the BH method: * *p* < 0.05; ** *p* < 0.01; *** *p* < 0.001. Experimental data and scatter plots of experimental versus predicted values associated with this analysis are provided in Fig. S1.

### Optimizing activin A and CHIR-99021 enables highly efficient, stable, and reproducible cardiogenic mesoderm induction

Given the positive effect of Act in the presence of suboptimal CHIR, we aimed to optimize the concentrations of these two factors for cardiogenic mesoderm induction. To achieve this, we applied response surface methodology (RSM) using a central composite design (CCD) (Fig. 2a) to model the induction of KDR^+^/PDGFR-α^+^ cells. CHIR has been reported to exhibit a steep dose response in cardiac induction [25], underscoring the need for careful optimization. Thus, we conducted two sequential RSM experiments (Table S2), where the concentration range in the second experiment was slightly shifted to better capture the response peak range. Additionally, data from both experiments were pooled to enhance statistical power and improve the accuracy of model coefficient estimation, following the procedure reported by Audet *et al.* [19].

**Fig. 2.**
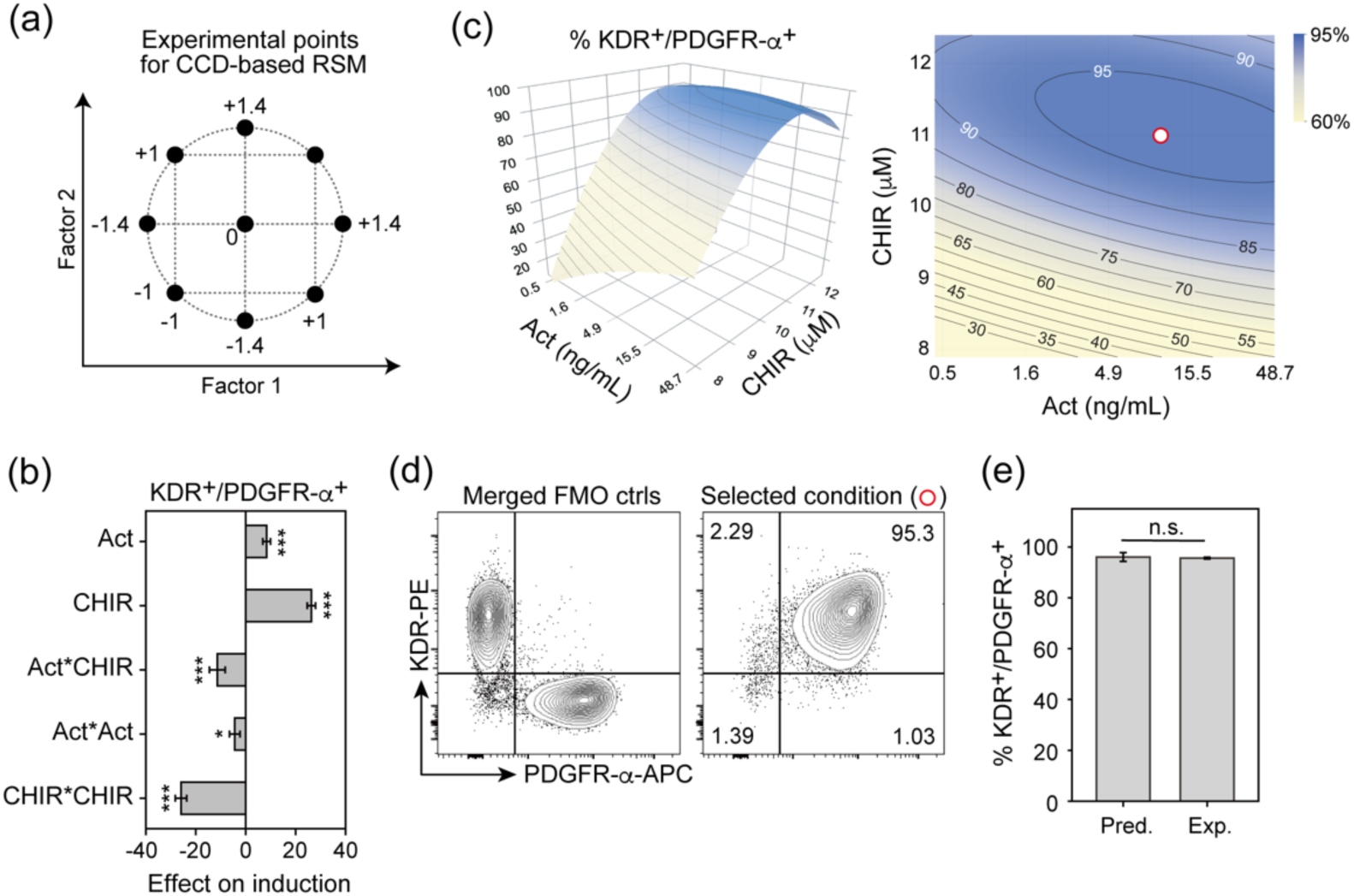
Factor optimization for cardiogenic mesoderm induction. (a) Geometric representation of experimental points in response surface methodology (RSM) based on a central composite design (CCD) with two factors. The scaled values range from −1.4 to +1.4, corresponding to minimum and maximum concentrations. The experiment was conducted twice using this design with slightly different concentration ranges specified in Table S2. (b) Results of the response surface analysis using the pooled data from the two experiments on the induction efficiency of the KDR^+^/PDGFR-α^+^ population. The effects are shown as the values of the corresponding coefficients in Equation (2), along with SEs. Statistical significance was assessed using an *F*-test, followed by *p*-value adjustment using the BH method: * *p* < 0.05; *** *p* < 0.001. Experimental data and a scatter plot of experimental versus predicted values associated with this analysis are provided in Fig. S2. (c) Response surface and contour plots illustrating the predicted induction efficiency of the KDR^+^/PDGFR-α^+^ population. The condition selected for subsequent experiments is marked by an open red circle. (b) Representative flow cytometry data on day 3 for the selected condition (right), with gating thresholds determined based on fluorescence-minus-one (FMO) controls (left). (c) Comparison of the predicted data (Pred.; *n* = 3 from the model) and experimental data on day 3 (Exp.; *n* = 3 independent experiments) for the selected condition. Data are presented as the mean ± SE. Statistical significance was assessed using a *t*-test. “n.s.” indicates not significant (*p* ≥ 0.05).

Response surface analysis showed negative quadratic effects for both factors (Fig. 2b), suggesting the presence of a response peak. The response surface and contour plots confirmed that our model successfully captured the peak region within the applied concentration range (Fig. 2c). We identified a parameter space predicting highly efficient induction of approximately 95%. Within this space, we selected the condition of 10 ng/mL Act and 11 μM CHIR (marked by an open red circle in Fig. 2c) as the candidate optimum, owing to its proximity to the space center and its practical usability. To validate this, we conducted repeated differentiation experiments using the selected condition, yielding a mean of 95.6% with a range from 95.1% to 96.3% across independent batches, which was comparable to the model’s prediction with no significant difference (Fig. 2d, e). Importantly, the experimental data indicated not only high efficiency but also minimal variability, demonstrating high stability and reproducibility.

Cardiogenic mesoderm is known to give rise to CMs, MCs, and ECs [15,34]. To confirm whether the cells induced under the selected condition represented true cardiogenic mesoderm with proper differentiation potential, they were stimulated with or without cardiovascular specification factors. As shown in Fig. S3, unstimulated cells or those treated with WNTi alone exclusively differentiated into MCs or CMs, respectively, both at >96%, consistent with the highly efficient induction of KDR^+^/PDGFR-α^+^ cells. Additionally, ECs emerged under VEGF stimulation. Thus, we successfully validated the selected condition for its ability to efficiently and reproducibly induce authentic cardiogenic mesoderm with proper differentiation potential.

### Multi-response modeling enables controlled trilineage co-differentiation

Based on the results that cardiogenic mesoderm, either unstimulated or stimulated with WNTi or VEGF, differentiated into MCs, CMs, or ECs (Fig. S3), we aimed to develop a controlled trilineage co-differentiation process by modulating these two stimuli. Given the importance of heterotypic cellular proportions in facilitating proper interactions, as observed in co-culture studies [10–14], achieving high controllability over the differentiation ratio was a key focus. To achieve this, we applied multi-response modeling for these trilineage cell emergence using CCD-based RSM, with WNTi and VEGF as process parameters (Fig. 3a).

**Fig. 3.**
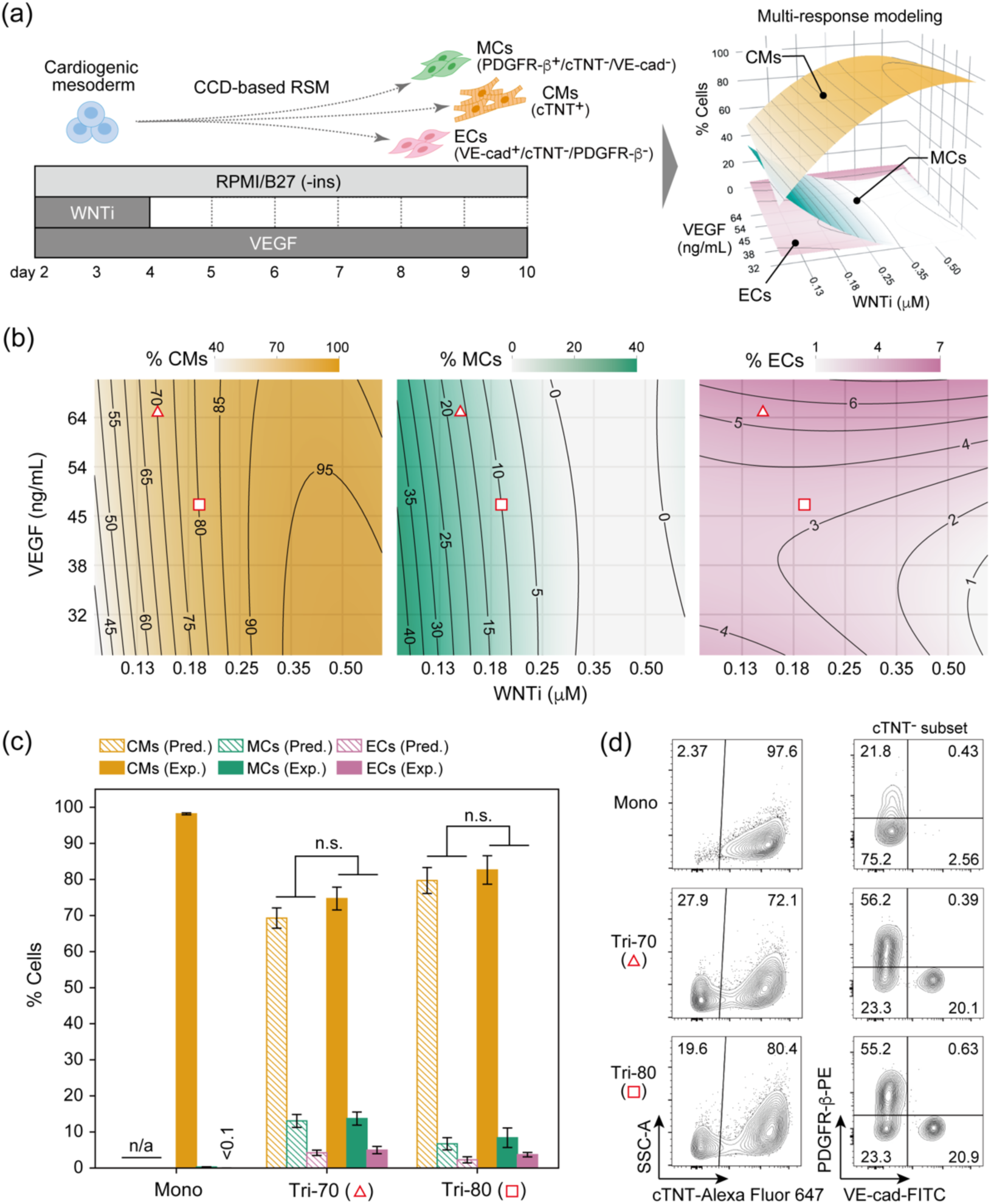
Controlled cardiac trilineage differentiation based on multi-response modeling. (a) Schematic representation of the experimental workflow for constructing multi-response models to predict co-differentiation ratios of CMs (cTNT+), MCs (PDGFR-β^+^/cTNT^−^/VE-cad^−^), and ECs (VE-cad^+^/cTNT^−^/ PDGFR-β^−^) using CCD-based RSM with WNTi and VEGF. These factors were introduced during the specified time windows at the concentrations outlined in Table S3. Experimental data, coefficient estimates, and scatter plots of experimental versus predicted values are provided in Fig. S4. (b) Contour plots illustrating the predicted induction efficiencies of CMs, MCs, and ECs. Two trilineage co-differentiation conditions, predicting 70% and 80% CM induction, labeled as Tri-70 (open red triangles; 0.14 μM WNTi and 65 ng/mL VEGF) and Tri-80 (open red squares; 0.19 μM WNTi and 47 ng/mL VEGF), were selected for subsequent experiments. (c) Comparison of predicted data (*n* = 5 from the models) and experimental data on day 10 (*n* = 5 independent experiments) for the selected conditions. Alongside trilineage differentiation, the monolineage differentiation (labeled as Mono), which exclusively generated CMs using 10 μM WNTi in an earlier experiment (Fig. S3), was also conducted (*n* = 5 independent experiments). Data are presented as the mean ± SE. Statistical significance was assessed using multivariate ANOVA. “n.s.” indicates not significant (*p* ≥ 0.05), and “n/a” indicates not available. (d) Representative flow cytometry data for Mono, Tri-70, and Tri-80 conditions on day 10.

As shown in Fig. 3b, multi-response modeling allowed the prediction of trilineage cell outcomes across the defined concentration range, facilitating the identification of potential conditions that steer co-differentiation toward the targeted proportions. To select the conditions, we leveraged findings from two recent studies. The first study confirmed that hiPSC-derived CM maturation was enhanced in co-culture with stromal cells and ECs at a CM proportion of 70% [13], providing insight into a preferable CM ratio for *in vitro* culture. Based on single-cell omics, the second study revealed an approximate 3:1 MC-to-EC ratio in the human heart ventricle [35]. Based on these findings, we selected a trilineage differentiation condition expected to generate CMs at 70% of the total cells, with the remaining cells exhibiting a 3:1 MC-to-EC ratio (labeled by open red triangles in Fig. 3b and referred to as the Tri-70 condition). Additionally, we selected a slightly different condition predicted to generate CMs at 80% of the total cells with a similar MC-to-EC ratio (labeled by open red squares in Fig. 3b and referred to as the Tri-80 condition). To validate the controllability of differentiation ratios, we conducted repeated differentiation experiments using Tri-70 and Tri-80 conditions alongside a monolineage differentiation condition (referred to as the Mono condition), which exclusively generated CMs under higher WNTi in an earlier experiment (Fig. S3). As shown in Fig. 3c, d, both Tri-70 and Tri-80 conditions generated CMs, MCs, and ECs, with their percentages comparable to the model’s predictions and showing no statistical difference, as confirmed using multivariate ANOVA. Thus, the results successfully confirmed the high controllability of our co-differentiation process in achieving targeted differentiation ratios.

### Enhanced maturation signature for cardiomyocytes from trilineage co-differentiation

To investigate gene expression differences between CMs derived from monolineage (Mono) and trilineage differentiation (Tri-70 and Tri-80), three-dimensional microtissues from each condition were generated on day 10 of differentiation using a microcavity plate (Fig. 4a). These microtissues were cultured for additional 30 days, with trilineage cultures supplemented with VEGF to support the vascular cell maintenance. qPCR analysis was then performed on day 40 (Fig. 4b). Among sarcomere genes, Tri-70 exhibited a significant upregulation of *MYH7* and a modest downregulation of *MYH6* compared with Mono, where *MYH7* and *MYH6* are the predominant myosin heavy chain isoforms in adult and fetal hearts, respectively. This led to a 10.4-fold increase in the *MYH7*/*MYH6* ratio. Furthermore, the *MHY7*/*MYH6* ratio in Tri-70 was 2.6-fold higher than in Tri-80. Similarly, the *MYL2*/*MYL7* ratio in Tri-70 was 5.0-fold higher than that in Mono, owing to an increase of *MYL2* and a decrease of *MYL7*, where *MYL2* is a mature ventricular marker, and *MYL7* represents an atrial but fetal marker. The *TNNI3*/*TNNI1* ratio, representing the shift from fetal to adult troponin isoforms, showed no statistically significant difference but an upward trend for Tri-70 due to a nearly 50% decrease in *TNNI1* expression compared with Mono. These results strongly support that CMs from Tri-70 exhibited the highest maturation level among the three conditions. While Tri-80 did not demonstrate significant differences from Mono in any of the three isoform ratios, all showed higher values, suggesting a modest enhancement of maturation. Thus, the overall results for sarcomere gene expression indicate that CMs derived from trilineage differentiation were more mature than those from monolineage differentiation. However, expression levels of calcium handling-related genes (*CACNA1C*, *RYR2*) and a β-oxidation-related gene (*FABP3*) remained unchanged, suggesting that any variable associated with trilineage differentiation and culture did not positively impact calcium-handing and metabolic maturation.

**Fig. 4.**
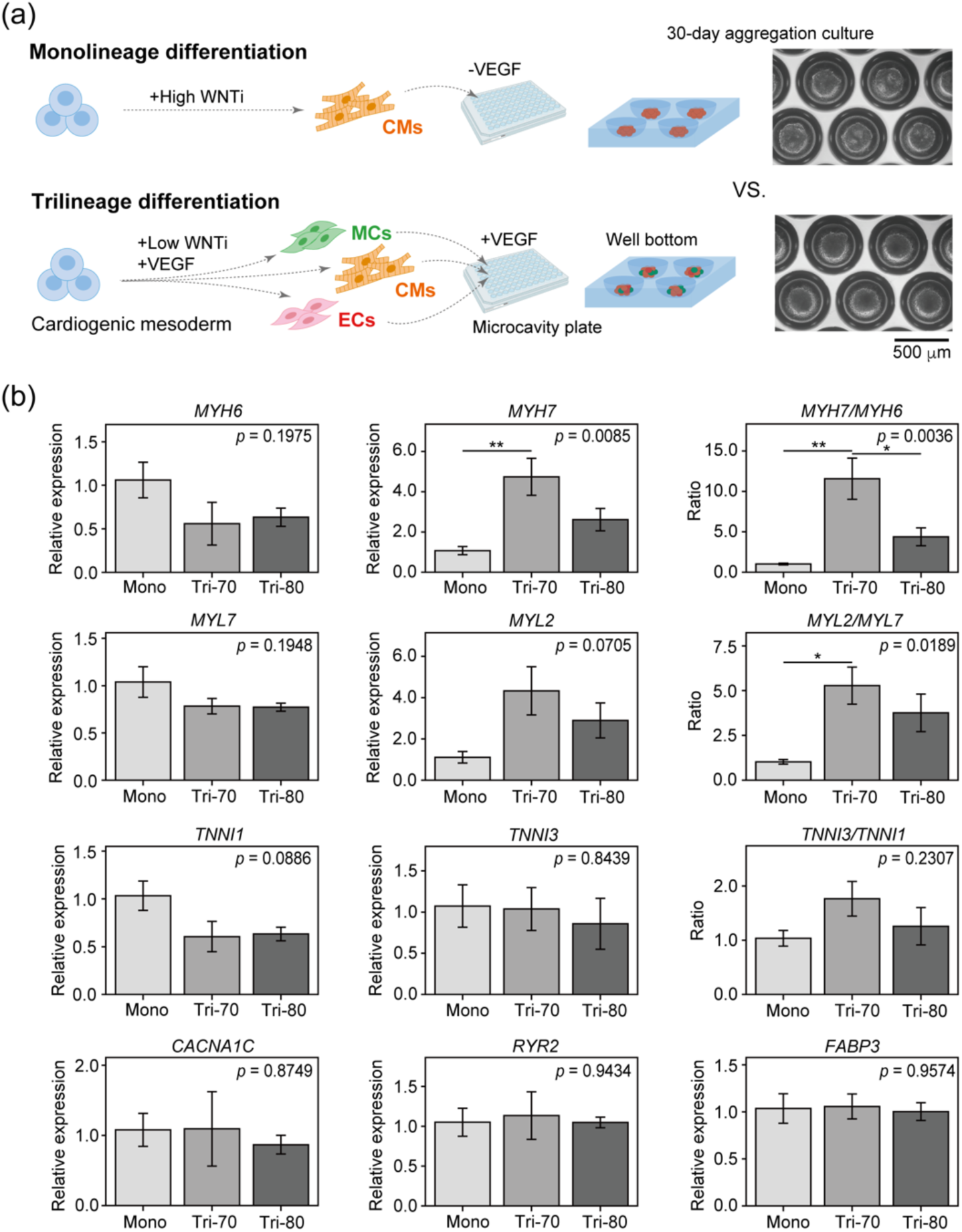
Comparison of cardiac maturation-associated gene expression between cells from mono and trilineage differentiation after prolonged culture. (a) Schematic representation of the experimental workflow for prolonged aggregation culture of cardiac cells derived from mono and trilineage differentiation. On day 10 of differentiation, cells from each culture were harvested, replated in a microcavity plate to form aggregates, and cultured for 30 days. Representative phase contrast images of cell aggregations on day 5 are shown, with the lower image displaying cell aggregates from the Tri-70 condition. (b) Comparison of gene expression levels for cells from Mono, Tri-70, and Tri-80 conditions using qPCR analysis after 30 days of aggregation culture. Relative expression is presented as the mean ± SE (*n* = 4 independent experiments). Overall statistical significance was assessed using one-way ANOVA, followed by Tukey’s post hoc test for pairwise comparisons. The overall *p*-value is displayed in the figure, and the results of Tukey’s test are indicated as follows: * *p* < 0.05; ** *p* < 0.01.

To further validate whether the observed sarcomere gene expression profile for cells from trilineage differentiation reflects a true signal of CM maturation, a single CM area, reflecting cell size, was measured after dissociation and replating. Cell size is a key hallmark for CM maturation linked to sarcomere development [36]. As shown in Fig. S5, CMs from Tri-70 and Tri-80 conditions exhibited a 1.4-fold and 1.3-fold increase in the average cell area, respectively, compared with those from the Mono conditions. The results support the differences in sarcomere gene expression profile and provide additional evidence supporting the relative maturation of CMs from trilineage differentiation, particularly in the Tri-70 condition, compared with monolinage differentiation.

## Discussion

We successfully developed a controlled cardiac multilineage co-differentiation process using the DoE approach. iPSCs were efficiently and stably differentiated into cardiogenic mesoderm and further differentiated into CMs, MCs, and ECs at targeted proportions based on multi-response modeling. Moreover, CMs from trilineage differentiation showed a more mature sarcomere gene expression profile than those from monolineage differentiation following prolonged culture.

For the evaluation of Act and BMP effects on cardiogenic mesoderm induction with CHIR (8–10 μM), Act showed a significant positive main effect, whereas no significant effects were observed for BMP (Fig. 1c). Our result for Act aligned with the previous report by Kim *et al*. [22]. However, Kim *et al*. have reported that BMP was inhibitory in the presence of optimized CHIR (12 μM), whereas Kadari *et al*. [29] have reported that BMP supplementation in the presence of CHIR (5 μM) promoted cardiac differentiation. Thus, the effect of BMP in the presence of CHIR remains inconsistent across the three studies. These discrepancies may be attributed to differences in WNT signal activity induced by CHIR treatment, given the required WNT activity for cardiac mesoderm induction and the signaling hierarchy during early development. WNT signaling governs differentiation fate, with low, intermediate, and high levels inducing endoderm, cardiac mesoderm, and paraxial/presomitic mesoderm, respectively [37]. This indicates the necessity of precisely modulating WNT activity to an intermediate level for cardiac differentiation. Moreover, BMP signaling activates the WNT pathway during gastrulation from hPSCs [38]. Based on these two prior findings, in a low CHIR condition, where WNT activity is insufficient, as in Kadari’s study (5 μM), BMP may serve as a positive enhancer of cardiac differentiation by increasing WNT activity. In contrast, BMP likely has little impact in a modest CHIR condition, where WNT activity is slightly below the optimal level as in our study (8–10 μM). However, in an optimized CHIR condition, as in Kim’s study (12 μM), BMP may have a negative effect due to excessive WNT activity. Thus, we believe that when both BMP and CHIR are used, the effect of BMP largely depends on the level of WNT signal activity induced by CHIR.

In the early stage of our process development, ensuring efficient, stable, and reproducible cardiogenic mesoderm induction was crucial, as it could significantly impact the overall process performance. The one-factor-at-a-time (OFAT) method, in which only one variable is adjusted while keeping all other variables constant, has been widely used in process development and various experimental studies. However, OFAT is prone to finding local optima and carries a significant risk of missing the true optimal condition [18] owing to its limited capacity to capture relationships between multiple parameters and outcomes. This is a critical yet often overlooked drawback. In this study, we employed sequential DoE experiments, consisting of FFD for factor evaluation and RSM for optimizing cardiogenic mesoderm induction. This systematic strategy yielded a highly efficient condition yielding KDR^+^/PDGFR-α^+^ cells at approximately 95% efficiency (Fig. 2d, e). As DoE reveals response behaviors within a defined multidimensional parameter space by statistical modeling, it has a lower risk of missing the true optimum. Thus, its application was a key to identifying this highly efficient condition. Cardiac differentiation is inherently prone to exhibit batch-to-batch variation [25,39,40], which is a common issue for stem cell differentiation. This variability may partly arise from insufficient optimization due to the limited ability of OFAT. Our optimized condition demonstrated minimal variability across independent experiments in this study, showing high stability and reproducibility. This is also supported by the high controllability of the subsequent co-differentiation (Fig. 3c), which would be unattainable with unstable initial differentiation. We believe that the high stability and reproducibility for cardiogenic mesoderm induction were also attributed to identifying true optimum through DoE, and that DoE serves as a powerful approach for engineering the stem cell differentiation process.

In developing the trilineage co-differentiation process, achieving high controllability over differentiation ratios is a central focus to facilitate proper heterotypic interactions, where controllability comprises two key aspects: accuracy and tunability. Accuracy refers to the extent to which the actual process outcome aligns with the targeted outcome, while tunability denotes the capability to adjust the process outcome to achieve desirable results by tuning parameters. In an earlier study, Masumoto *et al*. [15] pioneered the establishment of a co-differentiation process toward CMs, MCs, and ECs through VEGF stimulation alone for cardiac mesoderm. They have successfully demonstrated that transplanting cell sheets derived from the co-differentiation into rat infarcted hearts significantly improved cardiac function, demonstrating the potential application of the cardiac trilineage co-differentiation process in regenerative medicine. However, whether the experimentally observed differentiation proportion aligned with the desired outcome and to what extent the differentiation ratio was flexibly tunable by this single parameter remained uncertain, leaving the two key aspects of process controllability open for further exploration. In this study, we propose a new strategy to engineer a co-differentiation process based on multi-response modeling, which allowed us to predict differentiation ratios within a defined parameter space as well as identify potential conditions that steer the differentiation toward targeted proportions (Fig. 3b). The close match between targeted and actual differentiation ratios in the repeated validation runs (Fig. 3c, d) successfully demonstrated the high accuracy of our co-differentiation process. Furthermore, our strategy inherently possesses a high degree of tunability for differentiation ratio based on prediction without requiring additional experiments, facilitating process optimization to maximize the desired heterotypic interactions. Collectively, satisfying these two key aspects supports the high controllability of our process. Theoretically, our strategy can be extended to develop co-differentiation processes involving more cell types. Thus, our approach would be useful for engineering a co-differentiation process for regenerative medicine and drug development, where multicellular interactions are important.

In the gene expression analysis, CMs derived from trilineage co-differentiation exhibited maturation-associated sarcomere gene expression profile compared with those from monolineage differentiation (Fig. 4b). Enhanced maturation was further supported by increased CM size (Fig. S5). Since VEGF was supplemented only for cells from trilineage differentiation to support vascular cells during prolonged culture, a key question was whether the observed maturation signature resulted from heterotypic cellular interactions or merely a direct effect of VEGF on cardiomyocytes. To answer this, we conducted an additional experiment in which high-purity CMs were cultured with or without VEGF for a week and subjected to qPCR analysis. As shown in Fig. S6, VEGF showed no significant effect on sarcomere gene expression. The result strongly suggests that the enhanced sarcomere maturation signature was primarily driven by heterotypic interactions with non-myocytes, supporting the advantage of our co-differentiation process. Additionally, the maturation of cells from Tri-70 was more pronounced than those from Tri-80, indicating that the maturation level depends on the non-myocyte proportion. The results underscore the importance of tunability for differentiation ratios to maximize the impact of heterotypic interactions on CM properties and highlight the effectiveness of our process engineering strategy in making this possible. Although a detailed analysis of the mechanisms by which MCs and/or ECs modulate CM maturation is beyond the scope of this study, its exploration could provide valuable insights for future research.

Despite the positive signature for sarcomere maturation, we did not observe the upregulation of calcium handling and β-oxidation genes in cells from trilineage co-differentiation (Fig. 4b). The results suggest that the non-myocyte population under the present conditions cannot enhance broader aspects of CM maturation. One potential solution is to optimize the output cellular proportions. For example, a recent study has demonstrated that PSC-derived ECs can promote electrical CM maturation, when co-cultured at higher ratios than those in our study [10], suggesting that increasing EC proportions could be a promising approach. Yet, the predictive accuracy of our current models is limited to conditions with ECs at a maximum of approximately 7% (Fig. 3b) owing to the applied VEGF concentration range (<74 ng/mL) in the CCD experiment (Table S3). Thus, expanding the VEGF concentration range beyond 74 ng/mL in future improved models may enable the selection of conditions with a higher EC proportion, thereby enhancing CM-EC interactions to further promote CM maturation. Alternatively, another potential solution is to develop a new co-differentiation process that facilitates the co-emergence of an addition cell type with strong interactions with CMs. For instance, cardiac fibroblasts have been reported to arise from cardiac mesoderm in the presence of basic fibroblast growth factor [41] and to significantly drive CM maturation across multiple functional properties [13]. Therefore, a new process development with basic fibroblast growth factor as an additional parameter may further promote CM maturation by facilitating the co-emergence of cardiac fibroblasts and their interactions with CMs. Thus, our next step is to achieve broader CM maturation through these proposed process improvements.

## Conclusion

We proposed a new strategy to engineer a controlled cardiac multilineage differentiation process using the DoE approach. The key highlight was the high controllability of co-differentiation into CMs, MCs, and ECs at targeted proportions based on multi-response modeling. Additionally, gene expression analysis revealed that CMs from trilineage differentiation was a more matured sarcomere gene expression profile than those from monolineage differentiation. Thus, we believe that our strategy is useful for engineering stem cell co-differentiation processes for regenerative medicine and drug development where multicellular interactions are crucial. Nonetheless, broader aspects of CM maturation were not observed in the current process. To address this, our future research will focus on process improvement to achieve a more optimal differentiation ratio or to generate additional cell types, such as cardiac fibroblasts, that strongly enhance CM maturation.

## Supplementary information

**Supplementary data. Supplementary methods**; Evaluation of trilineage differentiation potential of cardiogenic mesoderm cells, cell size analysis for cardiomyocytes, and evaluation of the effect of VEGF addition on sarcomere gene expression. **Fig. S1.** Experimental data and scatter plots of experimental versus predicted values for factor evaluation in cardiogenic mesoderm induction. **Fig. S2.** Experimental data and a scatter plot of experimental versus predicted values for factor optimization in cardiogenic mesoderm induction. **Fig. S3.** Testing the differentiation potential of cardiogenic mesoderm cells toward CMs, MCs, and ECs. **Fig. S4.** Experimental data, coefficient estimates, and scatter plots of experimental versus predicted values for multi-response modeling of trilineage co-differentiation. **Fig. S5.** Comparison of single CM area after prolonged culture. **Fig. S6.** Evaluation of the effect of VEGF addition on sarcomere gene expression. **Table S1.** Design matrix and factor concentrations for a full factorial design experiment. **Table S2.** Design matrix and factor concentrations for a central composite design experiment with activin A and CHIR-99021. **Table S3.** Design matrix and factor concentrations for a central composite design experiment with WNTi and VEGF. **Table S4.** List of antibodies used in flow cytometry analysis. **Table S5.** List of primers used in this study.

## Abbreviations

iPSCs: Induced pluripotent stem cells
PSCs: Pluripotent stem cells
CMs: Cardiomyocytes
MCs: Mural cells
ECs: Endothelial cells
DoE: Design of experiments
FFD: Full factorial design
RSM: Response surface methodology
CCD: Central composite design
qPCR: Quantitative polymerase chain reaction
BH: Benjamini-Hochberg
ANOVA: Analysis of variance
RPMI/B27 (-ins): RPMI1640 supplemented with 2% B27 minus insulin
RPMI/B27 (+ins): RPMI1640 supplemented with 2% B27 containing insulin
CHIR: CHIR-99021
Act: Activin A
BMP: Bone morphogenetic protein-4
WNTi: WNT signaling inhibitor
KDR: Kinase insert domain receptor
PDGFR-α: Platelet-derived growth factor receptor-α
cTNT: Cardiac troponin T
PDGFR-β: Platelet-derived growth factor receptor-β
VE-cad: Vascular endothelial cadherin
OFAT: One-factor-at-a-time.

## Acknowledgements

We thank the Division for Medical Research Engineering at Nagoya University Graduate School of Medicine for providing access to the BD FACSCanto II flow cytometer and FlowJo software.

## Author contributions

Conceptualization: H.A., Y.K., and K.S.; Data curation: H.A. and Y.K.; Formal analysis: H.A. and Y.K.; Funding acquisition: H.A. and K.S.; Investigation: H.A. and Y.K.; Methodology: H.A. and Y.K.; Project administration: H.A.; Supervision: H.A., K.S. and H.H.; Validation: H.A. and Y.K.; Visualization: H.A. and Y.K.; Writing – original draft: H.A.; Writing – review & editing: H.A., K.S., and H.H. All authors read and approved the final manuscript.

## Availability of data and materials

The datasets used and/or analyzed during the current study are available from the corresponding author upon reasonable request.

## Declarations

### Ethics approval and consent to participate

Not applicable.

### Consent for publication

Not applicable.

## References

1. Brade T, Pane LS, Moretti A, Chien KR, Laugwitz K-L. Embryonic heart progenitors and cardiogenesis. Cold Spring Harb Perspect Med. 2013;3:a013847. https://pmc.ncbi.nlm.nih.gov/articles/PMC3784811/

2. Li Y, Du J, Deng S, Liu B, Jing X, Yan Y, et al. The molecular mechanisms of cardiac development and related diseases. Signal Transduct Target Ther. 2024;9:368. https://www.nature.com/articles/s41392-024-02069-8

3. Guo F, Zhang C-C, Yin X-H, Li T, Fang C-H, He X-B. Crosstalk between cardiomyocytes and noncardiomyocytes is essential to prevent cardiomyocyte apoptosis induced by proteasome inhibition. Cell Death Dis. 2020;11:783. https://www.nature.com/articles/s41419-020-03005-8

4. Tan S, Yang J, Hu S, Lei W. Cell-cell interactions in the heart: advanced cardiac models and omics technologies. Stem Cell Res Ther. 2024;15:362. https://stemcellres.biomedcentral.com/articles/10.1186/s13287-024-03982-z

5. Zhang H, Xue Y, Pan T, Zhu X, Chong H, Xu C, et al. Epicardial injection of allogeneic human-induced-pluripotent stem cell-derived cardiomyocytes in patients with advanced heart failure: protocol for a phase I/IIa dose-escalation clinical trial. BMJ Open. 2022;12:e056264. https://pubmed.ncbi.nlm.nih.gov/35523485/

6. Miyagawa S, Kainuma S, Kawamura T, Suzuki K, Ito Y, Iseoka H, et al. Case report: Transplantation of human induced pluripotent stem cell-derived cardiomyocyte patches for ischemic cardiomyopathy. Front Cardiovasc Med. 2022;9:950829. https://pubmed.ncbi.nlm.nih.gov/36051285/

7. Lee TYT, Coles JG, Maynes JT. iPSC-cardiomyocytes in the preclinical prediction of candidate pharmaceutical toxicity. Front Pharmacol. 2024;15:1308217. https://www.frontiersin.org/journals/pharmacology/articles/10.3389/fphar.2024.1308217/full

8. Wang EY, Rafatian N, Zhao Y, Lee A, Lai BFL, Lu RX, et al. Biowire model of interstitial and focal cardiac fibrosis. ACS Cent Sci. 2019;5:1146–58. https://pmc.ncbi.nlm.nih.gov/articles/PMC6661857/

9. Iseoka H, Miyagawa S, Sakai Y, Sawa Y. Cardiac fibrosis models using human induced pluripotent stem cell-derived cardiac tissues allow anti-fibrotic drug screening in vitro. Stem Cell Res. 2021;54:102420. 10.1016/j.scr.2021.102420

10. Lin Z, Garbern JC, Liu R, Li Q, Mancheño Juncosa E, Elwell HLT, et al. Tissue-embedded stretchable nanoelectronics reveal endothelial cell-mediated electrical maturation of human 3D cardiac microtissues. Sci Adv. 2023;9:eade8513. https://www.science.org/doi/10.1126/sciadv.ade8513

11. Lock RI, Graney PL, Tavakol DN, Nash TR, Kim Y, Sanchez E Jr, et al. Macrophages enhance contractile force in iPSC-derived human engineered cardiac tissue. Cell Rep. 2024;43:114302. https://pubmed.ncbi.nlm.nih.gov/38824644/

12. Voges HK, Foster SR, Reynolds L, Parker BL, Devilée L, Quaife-Ryan GA, et al. Vascular cells improve functionality of human cardiac organoids. Cell Rep. 2023;42:112322. https://www.cell.com/cell-reports/pdf/S2211-1247(23)00333-9.pdf

13. Giacomelli E, Meraviglia V, Campostrini G, Cochrane A, Cao X, van Helden RWJ, et al. Human-iPSC-derived cardiac stromal cells enhance maturation in 3D cardiac microtissues and reveal non-cardiomyocyte contributions to heart disease. Cell Stem Cell. 2020;26:862–879.e11. https://www.sciencedirect.com/science/article/pii/S1934590920302022

14. Masumoto H, Nakane T, Tinney JP, Yuan F, Ye F, Kowalski WJ, et al. The myocardial regenerative potential of three-dimensional engineered cardiac tissues composed of multiple human iPS cell-derived cardiovascular cell lineages. Sci Rep. 2016;6:29933. https://www.nature.com/articles/srep29933

15. Masumoto H, Ikuno T, Takeda M, Fukushima H, Marui A, Katayama S, et al. Human iPS cell-engineered cardiac tissue sheets with cardiomyocytes and vascular cells for cardiac regeneration. Sci Rep. 2014;4:6716. https://www.nature.com/articles/srep06716

16. Giacomelli E, Bellin M, Sala L, van Meer BJ, Tertoolen LGJ, Orlova VV, et al. Three-dimensional cardiac microtissues composed of cardiomyocytes and endothelial cells co-differentiated from human pluripotent stem cells. Development. 2017;144:1008–17. https://pubmed.ncbi.nlm.nih.gov/28279973/

17. Lipsitz YY, Timmins NE, Zandstra PW. Quality cell therapy manufacturing by design. Nat Biotechnol. 2016;34:393–400. https://www.nature.com/articles/nbt.3525

18. Bowden GD, Pichler BJ, Maurer A. A design of experiments (DoE) approach accelerates the optimization of copper-mediated 18F-fluorination reactions of arylstannanes. Sci Rep. 2019;9:11370. https://www.nature.com/articles/s41598-019-47846-6

19. Audet J, Miller CL, Eaves CJ, Piret JM. Common and distinct features of cytokine effects on hematopoietic stem and progenitor cells revealed by dose-response surface analysis. Biotechnol Bioeng. 2002;80:393–404. https://pubmed.ncbi.nlm.nih.gov/12325147/

20. Edgar JM, Michaels YS, Zandstra PW. Multi-objective optimization reveals time- and dose-dependent inflammatory cytokine-mediated regulation of human stem cell derived T-cell development. NPJ Regen Med. 2022;7:11. https://pubmed.ncbi.nlm.nih.gov/35087040/

21. Nakagawa M, Koyanagi M, Tanabe K, Takahashi K, Ichisaka T, Aoi T, et al. Generation of induced pluripotent stem cells without Myc from mouse and human fibroblasts. Nat Biotechnol. 2008;26:101–6. https://www.nature.com/articles/nbt1374

22. Kim M-S, Horst A, Blinka S, Stamm K, Mahnke D, Schuman J, et al. Activin-A and Bmp4 levels modulate cell type specification during CHIR-induced cardiomyogenesis. PLoS One. 2015;10:e0118670. https://pubmed.ncbi.nlm.nih.gov/25706534/

23. Lian X, Hsiao C, Wilson G, Zhu K, Hazeltine LB, Azarin SM, et al. Robust cardiomyocyte differentiation from human pluripotent stem cells via temporal modulation of canonical Wnt signaling. Proc Natl Acad Sci U S A. 2012;109:E1848–57. https://www.pnas.org/doi/10.1073/pnas.1200250109

24. Burridge PW, Matsa E, Shukla P, Lin ZC, Churko JM, Ebert AD, et al. Chemically defined generation of human cardiomyocytes. Nat Methods. 2014;11:855–60. https://pmc.ncbi.nlm.nih.gov/articles/PMC4169698/

25. Laco F, Woo TL, Zhong Q, Szmyd R, Ting S, Khan FJ, et al. Unraveling the inconsistencies of cardiac differentiation efficiency induced by the GSK3β inhibitor CHIR99021 in human pluripotent stem cells. Stem Cell Reports. 2018;10:1851–66. https://pubmed.ncbi.nlm.nih.gov/29706502/

26. Lee JH, Protze SI, Laksman Z, Backx PH, Keller GM. Human pluripotent stem cell-derived atrial and ventricular cardiomyocytes develop from distinct mesoderm populations. Cell Stem Cell. 2017;21:179–194.e4. https://pubmed.ncbi.nlm.nih.gov/28777944/

27. Kattman SJ, Witty AD, Gagliardi M, Dubois NC, Niapour M, Hotta A, et al. Stage-specific optimization of activin/nodal and BMP signaling promotes cardiac differentiation of mouse and human pluripotent stem cell lines. Cell Stem Cell. 2011;8:228–40. https://pubmed.ncbi.nlm.nih.gov/21295278/

28. Yang D, Gomez-Garcia J, Funakoshi S, Tran T, Fernandes I, Bader GD, et al. Modeling human multi-lineage heart field development with pluripotent stem cells. Cell Stem Cell. 2022;29:1382–1401.e8. https://pubmed.ncbi.nlm.nih.gov/36055193/

29. Kadari A, Mekala S, Wagner N, Malan D, Köth J, Doll K, et al. Robust generation of cardiomyocytes from human iPS cells requires precise modulation of BMP and WNT signaling. Stem Cell Rev. 2015;11:560–9. https://pubmed.ncbi.nlm.nih.gov/25392050/

30. Tohyama S, Fujita J, Fujita C, Yamaguchi M, Kanaami S, Ohno R, et al. Efficient large-scale 2D culture system for human induced pluripotent stem cells and differentiated cardiomyocytes. Stem Cell Reports. 2017;9:1406–14. https://pubmed.ncbi.nlm.nih.gov/28988990/

31. Miwa T, Idiris A, Kumagai H. A novel cardiac differentiation method of a large number and uniformly-sized spheroids using microfabricated culture vessels. Regen Ther. 2020;15:18–26. https://www.sciencedirect.com/science/article/pii/S2352320420300298

32. Fukushima H, Yoshioka M, Kawatou M, López-Dávila V, Takeda M, Kanda Y, et al. Specific induction and long-term maintenance of high purity ventricular cardiomyocytes from human induced pluripotent stem cells. PLoS One. 2020;15:e0241287. https://pubmed.ncbi.nlm.nih.gov/33137106/

33. Yang L, Soonpaa MH, Adler ED, Roepke TK, Kattman SJ, Kennedy M, et al. Human cardiovascular progenitor cells develop from a KDR+ embryonic-stem-cell-derived population. Nature. 2008;453:524–8. https://pubmed.ncbi.nlm.nih.gov/18432194/

34. Wang L, Zhang F, Duan F, Huang R, Chen X, Ming J, et al. Homozygous MESP1 knock-in reporter hESCs facilitated cardiovascular cell differentiation and myocardial infarction repair. Theranostics. 2020;10:6898–914. https://pubmed.ncbi.nlm.nih.gov/32550911/

35. Litviňuková M, Talavera-López C, Maatz H, Reichart D, Worth CL, Lindberg EL, et al. Cells of the adult human heart. Nature. 2020;588:466–72. https://pubmed.ncbi.nlm.nih.gov/32971526/

36. Guo Y, Cao Y, Jardin BD, Sethi I, Ma Q, Moghadaszadeh B, et al. Sarcomeres regulate murine cardiomyocyte maturation through MRTF-SRF signaling. Proc Natl Acad Sci U S A. 2021;118:e2008861118. https://www.pnas.org/doi/10.1073/pnas.2008861118

37. Zhao M, Tang Y, Zhou Y, Zhang J. Deciphering Role of Wnt Signalling in Cardiac Mesoderm and Cardiomyocyte Differentiation from Human iPSCs: Four-dimensional control of Wnt pathway for hiPSC-CMs differentiation. Sci Rep. 2019;9:19389. https://pubmed.ncbi.nlm.nih.gov/31852937/

38. Martyn I, Kanno TY, Ruzo A, Siggia ED, Brivanlou AH. Self-organization of a human organizer by combined Wnt and Nodal signalling. Nature. 2018;558:132–5. https://www.nature.com/articles/s41586-018-0150-y

39. Prondzynski M, Berkson P, Trembley MA, Tharani Y, Shani K, Bortolin RH, et al. Efficient and reproducible generation of human iPSC-derived cardiomyocytes and cardiac organoids in stirred suspension systems. Nat Commun. 2024;15:5929. https://pmc.ncbi.nlm.nih.gov/articles/PMC11251028/

40. Yang X, Chen D, Sun Q, Wang Y, Xia Y, Yang J, et al. A live-cell image-based machine learning strategy for reducing variability in PSC differentiation systems. Cell Discov. 2023;9:53. https://www.nature.com/articles/s41421-023-00543-1

41. Zhang J, Tao R, Campbell KF, Carvalho JL, Ruiz EC, Kim GC, et al. Functional cardiac fibroblasts derived from human pluripotent stem cells via second heart field progenitors. Nat Commun. 2019;10:2238. https://www.nature.com/articles/s41467-019-09831-5

