## supplementary data for "Engineering a Controlled Cardiac Multilineage Co-Differentiation Process Using Statistical Design of Experiments"

### Supplementary methods

#### Evaluation of trilineage differentiation potential of cardiogenic mesoderm cells

Differentiation was initiated under optimized CHIR and Act concentrations. On day 2, the medium was replaced with RPMI/B27 (-ins) containing no supplements, 10 μM WNTi, 50 ng/mL VEGF, or a combination of both. From day 4 onward, VEGF was retained, while WNTi was excluded, with media exchanged every 2–3 days. On day 10, the cells were harvested for flow cytometry analysis using the procedure outlined in the Methods.

#### Cell size analysis for cardiomyocytes

On day 40 of differentiation, equivalent to day 30 of aggregation culture, cell aggregates from mono and trilineage differentiation were collected and dissociated using the AscleStem Cardiomyocyte Dissociation Solution. The cells were then filtered through a 40 μm pore-size strainer (Corning, Cat. # 352340) and seeded into a glass chamber slide (Watson, Tokyo, Japan, Cat. # 192-008) pre-coated with Geltrex (1:50 dilution) at 10,000–22,000 cells/cm$^2$ in α-MEM supplemented with 10% fetal bovine serum, 1% penicillin-streptomycin, and 10 μM Y-27632. After 48 h, the medium was replaced with RPMI/B27 (+ins). After 4–6 days of seeding, once cells had sufficiently spread, they were fixed with 4% paraformaldehyde phosphate buffer solution (Fujifilm Wako Pure Chemical, Cat. # 163-20145) for 20 min and permeabilized with 0.3% Triton X-100 (Sigma-Aldrich, Cat. # T8787) for 5 min at room temperature. Blocking was then performed using phosphate-buffered saline containing 10% goat serum (Vector Laboratories, Newark, CA, USA, Cat. # S-1000) and 0.01% Triton X-100 for 1 h at room temperature. The cells were then incubated with a mouse anti-α-actinin antibody (1:400 dilution; Sigma-Aldrich, Cat. # A7811) for 1 h at room temperature, followed by incubation with a CF488A-conjugated goat anti-mouse IgG (1:1000 dilution; Biotium, Fremont, CA, USA, Cat. # 20018) and 1 μg/mL 4′,6-diamidino-2-phenylindole (Dojindo Laboratories, Kumamoto, Japan, Cat. # D523) for 1 h at room temperature. Stained images were captured using a BZ-X700 fluorescence microscope (Keyence, Osaka, Japan). Individual α-actinin-positive cell area was measured using the ImageJ software (version 1.54f; National Institutes of Health, Bethesda, MD, USA).

#### Evaluation of the effect of VEGF addition on sarcomere gene expression

Differentiation was performed with 10 μM WNTi from days 2 to 4, as described earlier, to exclusively generate CMs. On day 10, the cells were harvested and dissociated using TrypLE Express. The cells were then suspended in α-MEM supplemented with 10% fetal bovine serum, 1% penicillin-streptomycin, and 10 μM Y-27632, with or without 50 ng/mL VEGF, and seeded into a 48-well plate (AGC Techno Glass, Shizuoka, Japan, Cat. # 3830-048) pre-coated with Geltrex (1:50 dilution) at 10,000 cells/cm$^2$. After two days of culture, the medium was replaced with RPMI/B27 (+ins), with or without 50 ng/mL VEGF, and cultured for an additional five days, ensuring that CMs in one of the two conditions were exposed to VEGF for a total of seven days. Finally, qPCR analysis was performed using the procedure outlined in the Methods.

### Supplementary figures

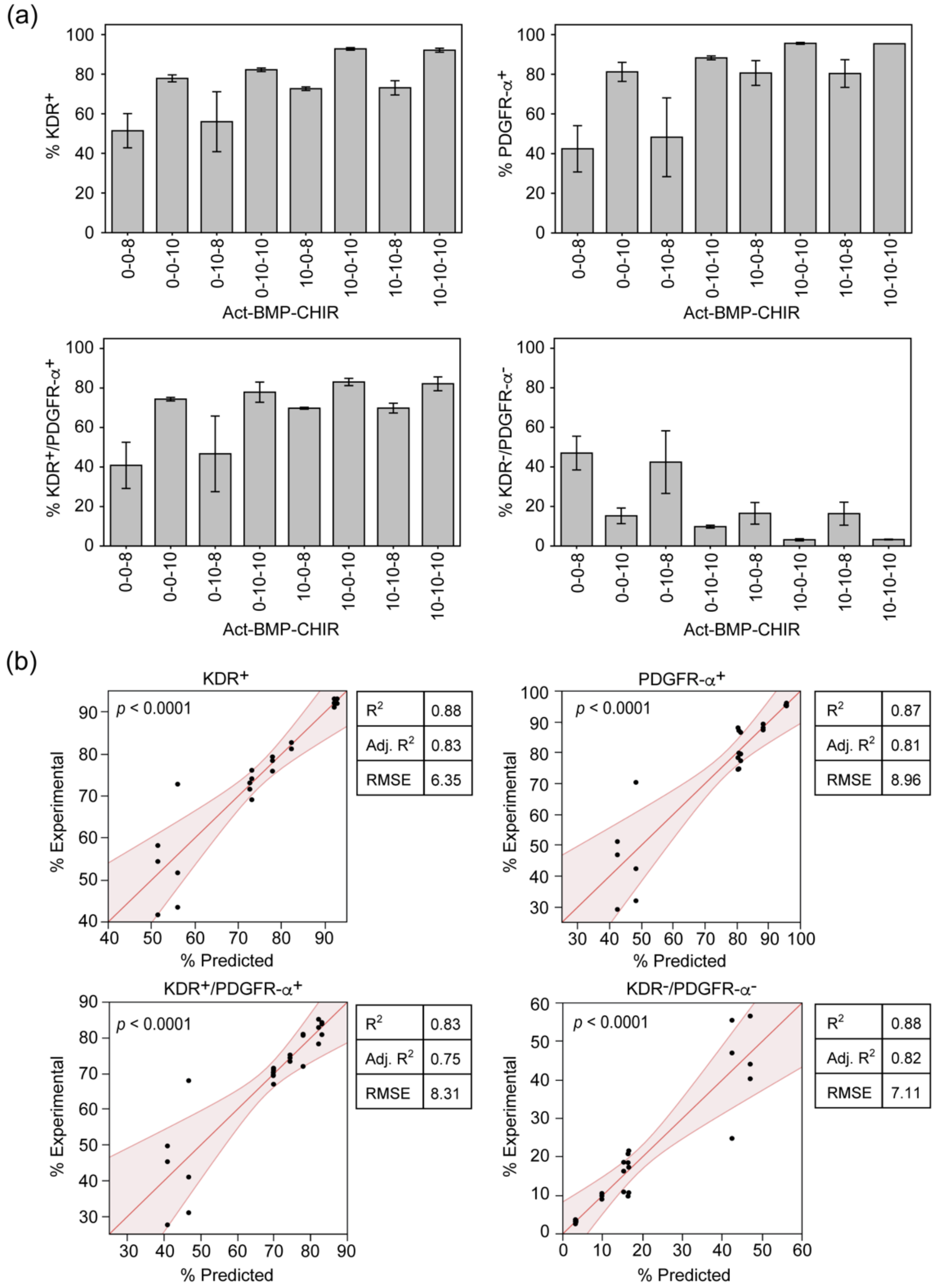


**Fig. S1. Experimental data and scatter plots of experimental versus predicted values for factor evaluation in cardiogenic mesoderm induction.** (a) Percentages of the cell populations of interest in the FFD experiment. Conditions are represented as combinations of the three-factor concentrations: Act (ng/mL), BMP (ng/mL), and CHIR (μM), in that order. Data are presented as the mean ± standard deviation (SD), with $n = 3$. (b) Scatter plots of experimental versus predicted values. The light red zone represents the 95% confidence region. The overall significance of the regression model was evaluated using ANOVA. The resultant *p*-value is displayed within each figure. A summary of the model fit is shown on the right side of each plot.

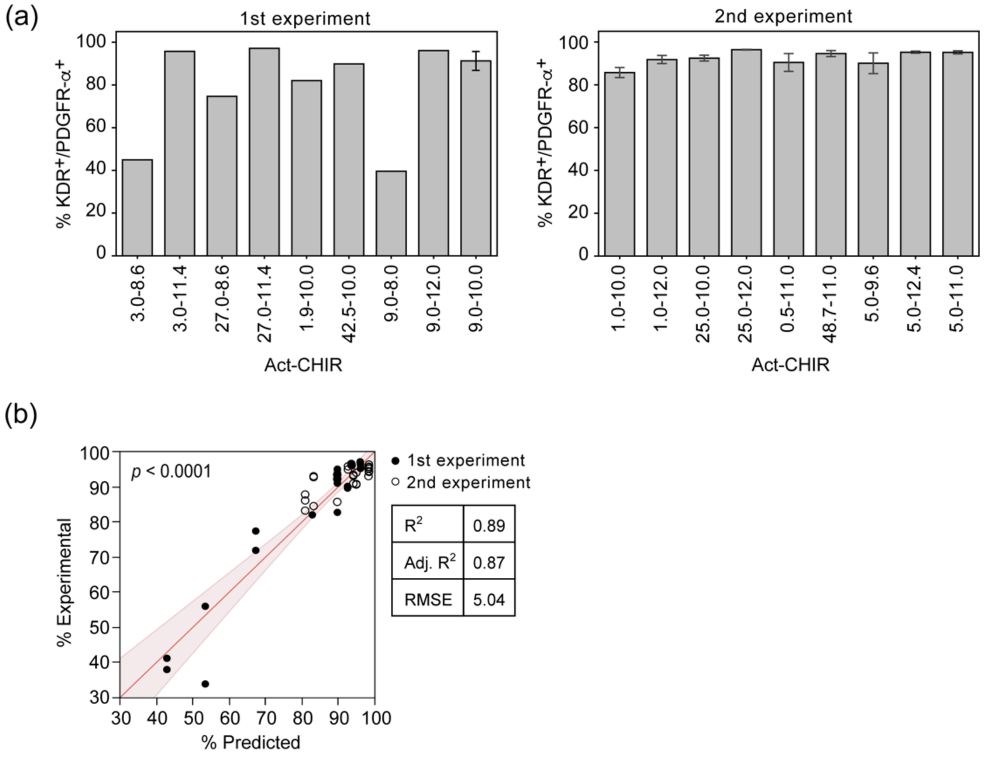


**Fig. S2. Experimental data and a scatter plot of experimental versus predicted values for factor optimization in cardiogenic mesoderm induction.** (a) Percentages of the $KDR^+$/PDGFR-$\alpha^+$ population in the two CCD experiments. Conditions are represented as combinations of the two-factor concentrations, Act (ng/mL) and CHIR (μM), in that order. In the first experiment, the center point condition labeled as "9.0-10.0" was measured with $n = 6$, while all other conditions with $n = 2$. In the second experiment, the center point condition labeled as "5.0-11.0" was measured with $n = 9$, while all other conditions with $n = 3$. Data are presented as the mean for the conditions with $n < 3$ and as the mean ± SD for conditions with $n \geq 3$. (b) A scatter plot of experimental versus predicted values. The data points from the first and second experiments are displayed by filled and open circles, respectively. The light red zone represents the 95% confidence region. The overall significance of the regression model was evaluated using ANOVA. The resultant $p$-value is displayed within the figure. A summary of the model fit is shown on the right side of the plot.

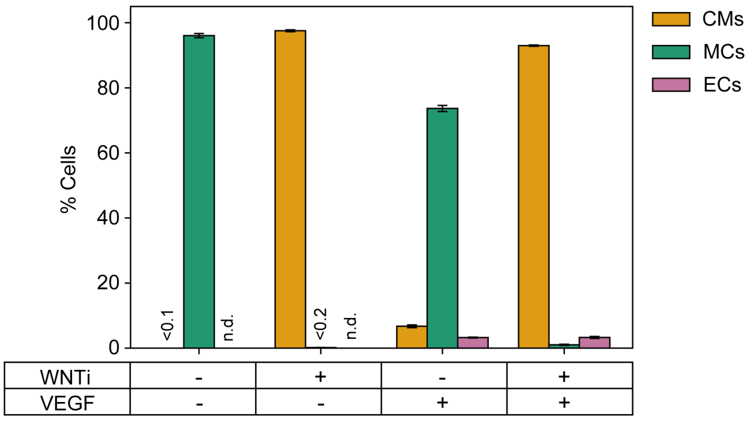


**Fig. S3. Testing the differentiation potential of cardiogenic mesoderm cells toward CMs, MCs, and ECs.** From day 2 of differentiation, cells were cultured under four conditions based on the presence (+) or absence (-) of WNTi and VEGF. The time window for the addition of each factor is shown in Fig. 3a. On day 10, cells were harvested and analyzed using flow cytometry to determine the percentages of CMs (cTNT$^{+}$), MCs (PDGFR-β$^{+}$/cTNT$^{-}$/VE-cad$^{-}$), and ECs (VE-cad$^{+}$/cTNT$^{-}$/ PDGFR-β$^{-}$). Data are presented as the mean ± SD ($n$ = 3). "n.d." indicates not detected.

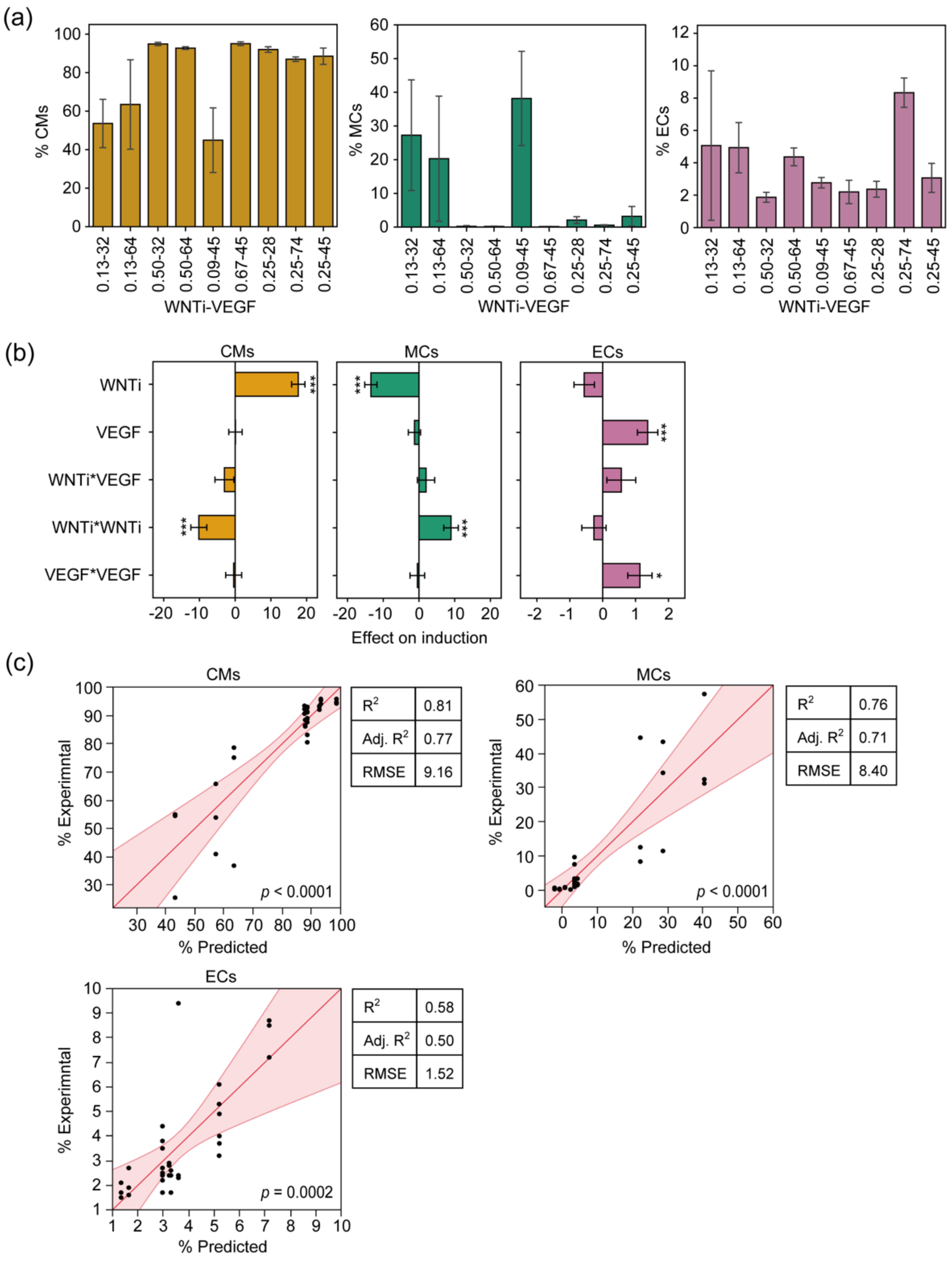


**Fig. S4. Experimental data, coefficient estimates, and scatter plots of experimental versus predicted values for multi-response modeling of trilineage co-differentiation.** (a) Percentages of trilineage cells in the CCD experiment. Conditions are represented as combinations of the two-factor concentrations, WNTi (μM) and VEGF (ng/mL), in that order. Data are presented as mean ± SD, with $n = 9$ for the center point condition labeled as "0.25-45" and $n = 3$ for all other conditions. (b) Results of the response surface analysis. The effects are shown as the values of the corresponding coefficients in Equation (2), along with SEs. Statistical significance was assessed using an $F$-test, followed by $p$-value adjustment using the BH method: * $p < 0.05$; *** $p < 0.001$. (c) Scatter plots of experimental versus predicted values. The light red zone represents the 95% confidence region. The overall significance of the regression model was evaluated using ANOVA. The resultant $p$-value is displayed within each figure. A summary of the model fit is shown on the right side of each plot.

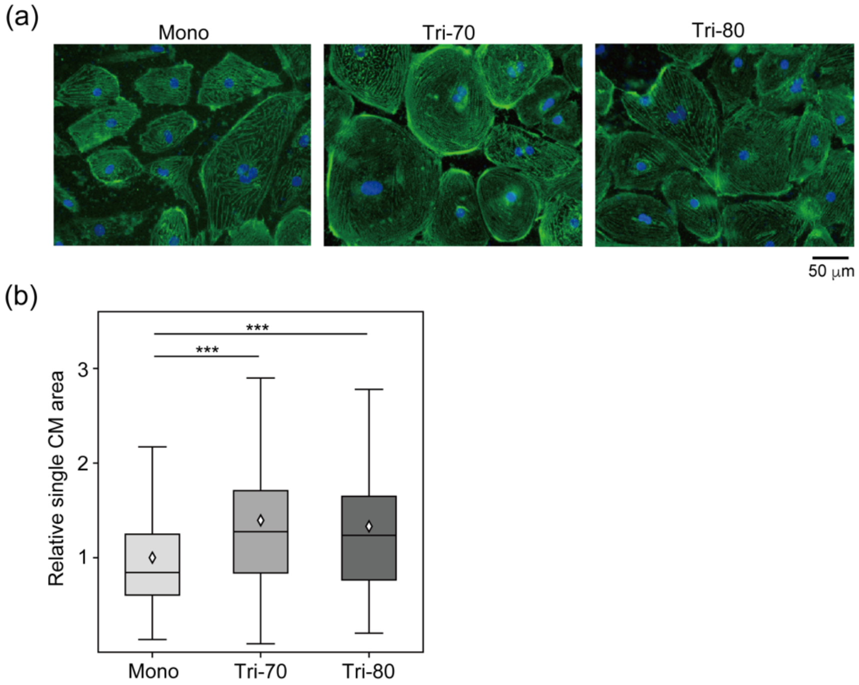


**Fig. S5. Comparison of single CM area after prolonged culture.** (a) Representative fluorescent microscopy images of replated cells stained for sarcomeric α-actinin (green) and nuclei (blue). Cells derived from Mono, Tri-70, and Tri-80 conditions were dissociated after 30 days of prolonged aggregation culture, then replated and subjected to fluorescent staining. (b) Relative single CM area. Data were collected from α-actinin-positive cells and pooled from two independent experiments ($n$ = 63–90). The mean value is indicated by an open diamond. Overall statistical significance was assessed using the Kruskal-Wallis test, followed by Dunn's post hoc test for pairwise comparisons: *** $p < 0.001$.

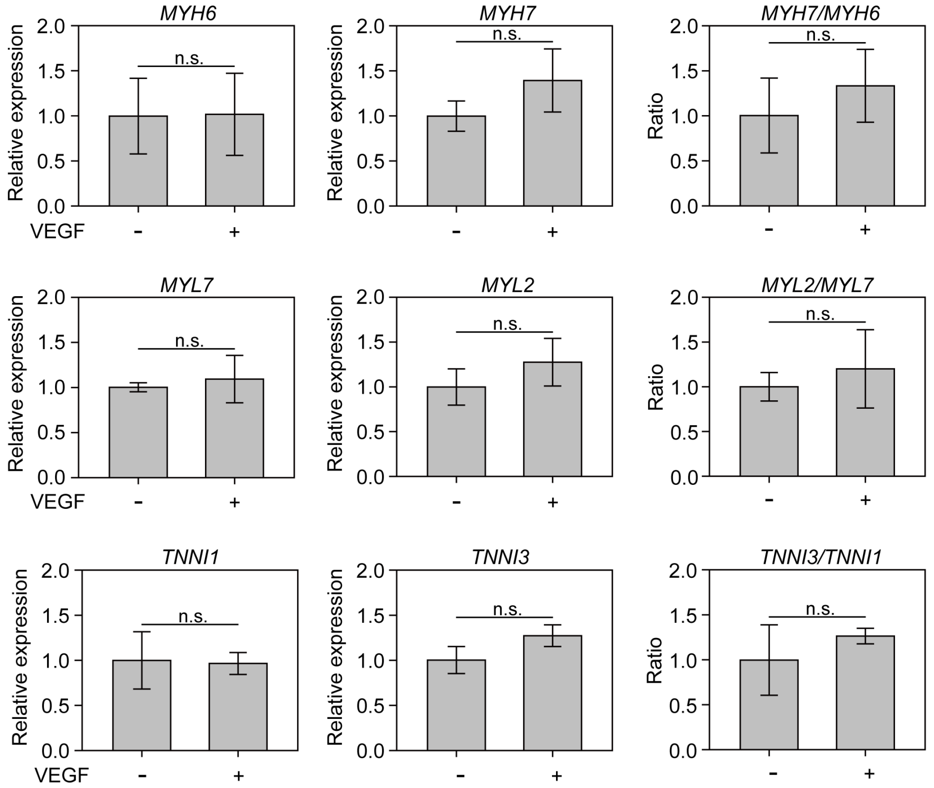


**Fig. S6. Evaluation of the effect of VEGF addition on sarcomere gene expression.** CMs were exclusively induced using the monolineage differentiation protocol. On day 10, CMs were harvested, replated, and cultured for seven days in the presence or absence of VEGF (50 ng/mL), followed by qPCR analysis. The percentage of CMs used in this experiment was 94.7%, determined using flow cytometry on day 10 of differentiation. Data are presented as the mean ± SD ($n$ = 3). Statistical significance was assessed using a $t$-test. "n.s." indicates not significant ($p \geq 0.05$).

### Supplementary Tables

**Table S1. Design matrix and factor concentrations for a full factorial design experiment**

| Scaled value | | | Concentration | | |
|---|---|---|---|---|---|
| Act | BMP | CHIR | Act (ng/mL) | BMP (ng/mL) | CHIR (μM) |
| -1 | -1 | -1 | 0 | 0 | 8 |
| -1 | -1 | +1 | 0 | 0 | 10 |
| -1 | +1 | -1 | 0 | 10 | 8 |
| -1 | +1 | +1 | 0 | 10 | 10 |
| +1 | -1 | -1 | 10 | 0 | 8 |
| +1 | -1 | +1 | 10 | 0 | 10 |
| +1 | +1 | -1 | 10 | 10 | 8 |
| +1 | +1 | +1 | 10 | 10 | 10 |

Each experimental point was measured with $n = 3$.

**Table S2. Design matrix and factor concentrations for a central composite design experiment with activin A and CHIR-99021**

| Scaled value | | Concentration | | | |
|---|---|---|---|---|---|
| | | 1st experiment | | 2nd experiment | |
| Act | CHIR | Act (ng/mL) | CHIR (μM) | Act (ng/mL) | CHIR (μM) |
| -1 | -1 | 3.0 | 8.6 | 1.0 | 10.0 |
| -1 | +1 | 3.0 | 11.4 | 1.0 | 12.0 |
| +1 | -1 | 27.0 | 8.6 | 25.0 | 10.0 |
| +1 | +1 | 27.0 | 11.4 | 25.0 | 12.0 |
| -1.414 | 0 | 1.9 | 10.0 | 0.5 | 11.0 |
| +1.414 | 0 | 42.5 | 10.0 | 48.7 | 11.0 |
| 0 | -1.414 | 9.0 | 8.0 | 5.0 | 9.6 |
| 0 | +1.414 | 9.0 | 12.0 | 5.0 | 12.4 |
| 0 | 0 | 9.0 | 10.0 | 5.0 | 11.0 |

Each experimental point, except for the center point where both scaled values are 0, was measured with $n = 2$ in the first experiment and $n = 3$ in the second experiment. The center point was measured with $n = 6$ in the first experiment and $n = 9$ in the second experiment. Factor concentration ($c$) is defined as follows: $c = c_0 \times 3^X$ for Act and $c = c_0 + 1.4 \times X$ for CHIR in the first experiment, and $c = c_0 \times 5^X$ for Act and $c = c_0 + X$ for CHIR in the second experiment, where $X$ is the scaled value and $c_0$ is the concentration at $X = 0$.

**Table S3. Design matrix and factor concentrations for a central composite design experiment with WNTi and VEGF**

| Scaled value | | Concentration | |
|---|---|---|---|
| WNTi | VEGF | WNTi (μM) | VEGF (ng/mL) |
| -1 | -1 | 0.13 | 32 |
| -1 | +1 | 0.13 | 64 |
| +1 | -1 | 0.50 | 32 |
| +1 | +1 | 0.50 | 64 |
| -1.414 | 0 | 0.09 | 45 |
| +1.414 | 0 | 0.67 | 45 |
| 0 | -1.414 | 0.25 | 28 |
| 0 | +1.414 | 0.25 | 74 |
| 0 | 0 | 0.25 | 45 |

Each experimental point, except for the center point where both scaled values are 0, was measured with $n = 3$. The center point was measured with $n = 9$. Factor concentration ($c$) is defined as follows: $c = c_0 \times 2^X$ for WNTi and $c = c_0 \times 1.41^X$ for VEGF, where $X$ is the scaled value and $c_0$ is the concentration at $X = 0$.

**Table S4. List of antibodies used in flow cytometry analysis**

| Antigen | Species | Vender | Clone | Cat. # | Fluorophore | Dilution |
|---|---|---|---|---|---|---|
| KDR | Mouse | BioLegend | A16085H | 393003 | PE | 1:50 |
| PDGFR-α | Mouse | BioLegend | 16A1 | 323511 | APC | 1:50 |
| cTNT | Mouse | BD Biosciences | 13-11 | 565744 | Alexa Fluor 647 | 1:50 |
| PDGFR-β | Rabbit | Sino Biological | 206 | 10514-R206-P | PE | 1:50 |
| VE-cad | Rabbit | Sino Biological | 048 | 10433-R048-F | FITC | 1:50 |

**Table S5. List of primers used in this study**

| Gene name | | 5′–3′ primer sequence | $T_m$ (°C) [a] | Product size (bp) | Accession # [b] |
|---|---|---|---|---|---|
| RPL37A | F | GTGGTTCCTGCATGAAGACAGTG | 61.9 | 84 | NM_000998.5 |
| | R | TTCTGATGGCGGACTTTACCG | 60.4 | | |
| TNNI1 | F | GTGGGTGACTGGAGGAAGAA | 58.9 | 156 | NM_003281.4 |
| | R | GTGAGCTGGGTTGGAGAAGA | 59.3 | | |
| TNNI3 | F | CACCTCAAGCAGGTGAAGAAG | 58.9 | 129 | NM_000363.5 |
| | R | CAGGAAGGCTCAGCTCTCAA | 59.4 | | |
| MYL7 | F | CCTGAGTGCCTTCCGCATGT | 62.8 | 133 | NM_021223.3 |
| | R | GGGTGTCAGGGCGAACATCT | 62.2 | | |
| MYL2 | F | CCCTGACGTGACTGGCAACT | 62.4 | 164 | NM_000432.4 |
| | R | AGGTAGGGACAGAGGCGGTA | 61.6 | | |
| MYH6 | F | TGCGGCCCAGATTCTTCAGG | 62.5 | 154 | NM_002471.4 |
| | R | ATGTCAAAGGGCCGGGTCTG | 62.5 | | |
| MYH7 | F | AGCCAACACCAACCTGTCCA | 61.9 | 138 | NM_000257.4 |
| | R | TCATTCAAGCCCTTCGTGCC | 61.0 | | |
| RYR2 | F | TGAGTGCCCCAAGCATCTCG | 62.5 | 144 | NM_001035.3 |
| | R | CATGTCGCCCTCCAAGCAGA | 62.5 | | |
| CACNA1C | F | TCCGCTCTGCCTCACTAGGT | 62.5 | 112 | NM_199460.4 |
| | R | ACCAGATGCAAGGGCAGGAC | 62.5 | | |
| FABP3 | F | TGGGGGTGGAGTTCGATGAG | 61.3 | 125 | NM_001320996.2 |
| | R | CTCCCGCACAAGTGTGGTCT | 62.4 | | |

[a] The melting temperature ($T_m$) values were obtained using the NCBI Primer-BLAST tool.

[b] Accession numbers are provided for the representative isoform of each gene when multiple isoforms are present. Primers were designed to detect all isoforms classified as "REVIEWED" for each gene in the NCBI Gene database.
